# Scale-independent topological interactions drive the first fate decision in the *Drosophila* embryo

**DOI:** 10.1101/2023.10.11.561879

**Authors:** Woonyung Hur, Arghyadip Mukherjee, Luke Hayden, Ziqi Lu, Anna Chao, Noah P. Mitchell, Sebastian J. Streichan, Massimo Vergassola, Stefano Di Talia

## Abstract

During embryogenesis, the earliest cell fate decision is often linked to nuclear positioning, whose control arises from the integration of the cell cycle oscillator and associated cytoskeletal dynamics. Yet, the mechanisms that ensure that the correct number of nuclei move to the appropriate place remain poorly understood. Here, using light sheet microscopy, we show that in *Drosophila* embryos spindle orientation controls which nuclei migrate towards the cortex and which remains inside the embryo, thereby determining nuclear fate and the number of cells undergoing development. Combining computational methods inspired by integral geometry and manipulations of cell cycle genes, we show that spindle orientation is controlled by topological spindle-spindle interactions and not by internuclear distance. Using arguments describing the behavior of space-filling systems, we develop a theory for topological dependency in microtubule structures. Our work shows how topological interplay of microtubule mechanics can ensure robust control of density and cell fate determination.

## INTRODUCTION

Collective dynamics are ubiquitous in living systems, ranging from bacterial biofilms to animal behavior [51, 26, 43, 7]. Theoretical analyses of such dynamics have recently concentrated on the impact of metric (scale-dependent) *vs.* topological (scale-independent, but geometry-dependent) interactions [7]. It has been argued that topological interactions can contribute to the robustness of collective dynamics [4]. Yet, how such interactions arise from basic underlying processes is still unclear. Here, we address this question in the context of robustness in early embryogenesis, more specifically in nuclear migration in early *Drosophila* embryos. Through a combination of live imaging, computational image analysis and theory, we show how topological interactions can become dominant among microtubule structures and how such interactions contribute to the robustness of nuclear positioning in early *Drosophila* embryos.

Nuclear positioning is crucial for the proper functioning of cells and has been linked to several important cellular decisions [2, 27]. This is particularly relevant in early embryogenesis when, in species ranging from insects to mammals, the first fate decision is tightly linked to nuclear/cellular positioning [36, 9, 39]. In insect species where embryos develop as syncytial blastoderms, mechanisms of nuclear positioning ensure that hundreds of nuclei move to their correct position within the large embryo [19, 16, 20]. These mechanisms operate during the rapid cleavage divisions of early embryogenesis thus requiring fast and robust coordination of the activity of the cell cycle oscillator and cytoskeletal components [5]. The nature and dynamics of these mechanisms, as well as their robustness, remain to be elucidated. Geometrical constraints are likely to impact the emergence and maintenance of collective nuclear movements. Much like biochemical feedback mechanisms, the dynamic feedback between structure/geometry and the activity of cytoskeletal components possibly lies at the heart of robustness in morphogenesis [29, 10]. Here, we consider this feedback mechanism in the context of cortical migration of nuclei in the syncytial *Drosophila* embryo, a collective process driven by large-scale cytoskeletal dynamics.

The early development of *Drosophila melanogaster* is characterized by 13 rapid and synchronous cleavage divisions that result in a blastoderm containing between 5000-6000 nuclei on the surface and about a hundred yolk nuclei in the core of the embryo [22]. These cleavage divisions are accompanied by significant nuclear movements that ensure that nuclei are appropriately localized. Two main processes contribute to nuclear positioning: axial expansion and cortical migration. During nuclear cycles 4-6, nuclei spread along the anterior-posterior axis in a process, named axial expansion, that involves localized cell cycle oscillations, actomyosin driven cortical contractions and cytoplasmic flows [49, 59, 16]. Following this phase, in which nuclei rapidly move along the AP axis transported by large scale cytoplasmic flows [16, 37], the nuclei collectively migrate towards the cortex during cycles 7-9 [3, 24] (see Fig.1A). During cortical migration, the flows are significantly tamed and the motion of the nuclei is driven by microtubule structures, most likely asters and mitotic spindles [3, 24, 18]. During this process, a large fraction of nuclei move from the center of the embryo to the cortex while another smaller fraction remains inside [20]. In *Drosophila* embryos, about 70% of nuclei collectively migrate to the surface by cycle 10 (*∼* 350 out of 512), while the remaining 30% (*∼* 150 out of 512) stay in the middle/center and become yolk nuclei.

**Fig. 1.**
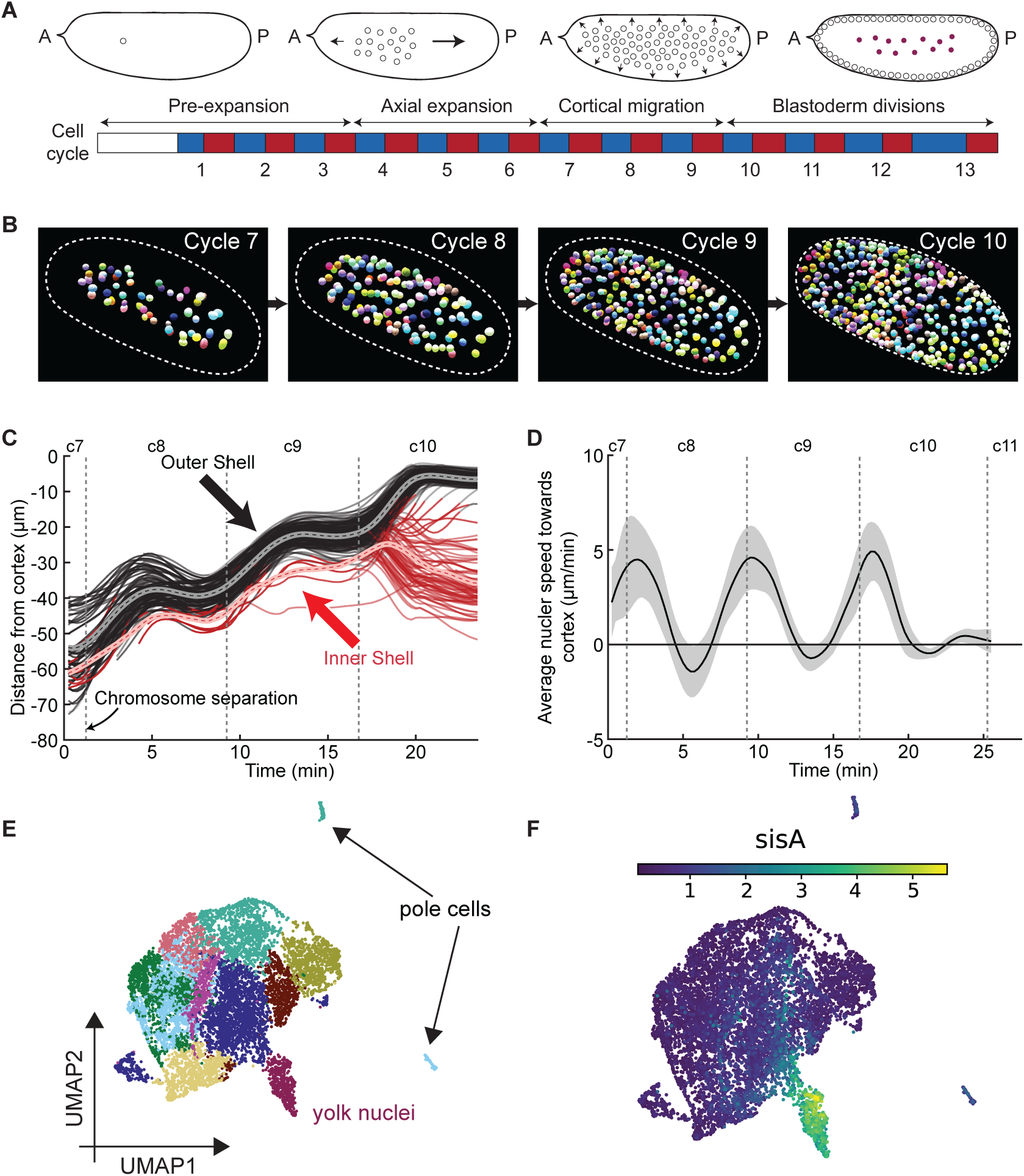
Symmetry breaking in nuclear positioning underlies embryo-yolk fate choice. **A,** Early development of the fruit fly *Drosophila melanogaster* embryo is schematised, marking morphogenetic events upto the 13th synchronous cell cycle. **B,** Segmented and tracked nuclei during the cortical migration phase are shown during the progression from cycle 7 to cycle 10. The white dashed line marks the final boundary at cycle 10. **C,** Distance of each nucleus from the embryo cortex is plotted against time. Events of chromosome separation are marked by dashed vertical lines. The bifurcated trajectories of outer and inner shells are indicated in black and red, respectively. Their bifurcation becomes particularly marked at the transition between cycles 9 and 10. **D,** Average nuclear speed is shown, in alignment with cell cycle phase. Solid line indicates the mean and the shaded region marks the standard deviation. **E,** An UMAP projection is shown for a single nuclei RNAseq data at the onset of maternal to zygotic transition, cluster with yolk nuclei is marked with dark brown. Reanalysed from [1]. **E,** The expression profile for the transcription factor SisA is shown to highlight its enrichment in the yolk nuclei cluster. Colorbar indicates relative enrichment in logarithmic scale.

Those nuclei that reach the cortex continue to proliferate and give rise to the cells undergoing morphogenesis and patterning of the embryo [22, 24]. Accurate control of nuclear density at the embryo surface plays a crucial role in the decision whether nuclei divide or not prior to the maternal-to-zygotic transition (MZT). This decision is made collectively by nuclei [28]. Yet, fluctuations of about 20-30% in local nuclear density could be sufficient to generate a variable number of divisions in different regions of the embryo [38]. Unlike the blastoderm nuclei, the yolk nuclei stop proliferating around cycle 10 and become polyploid [48]. Yolk nuclei have been implicated in the control of several developmental processes. Specifically, they drive the expression of several genes important for embryonic development. These genes include transcription factors (Sisterless A and Serpent), important for sex determination [21], endoderm, midgut and humoral immune system specification [62, 57], and a cytochrome P450 (Spook) involved in the synthesis of the insect hormone ecdysone, which plays a range of functions in *Drosophila* development [45]. Thus, the proper control of cortical migration is essential to establish that both the proper number of cells and the proper number of yolk nuclei are specified prior to the MZT and gastrulation. In spite of the crucial role of this morphogenetic process in embryogenesis, the mechanisms that control cortical migration, and more specifically the nuclear fate decision separating blastoderm and yolk nuclei, remain poorly understood.

Previous experiments in fixed embryos have shown that nuclear movements are limited to mitotic exit/early interphase and estimated the migration period to be about 30 seconds in anaphase/telophase [24]. Since this time period roughly coincide with the time of chromosome separation, the mitotic spindle was proposed to be the main driver of cortical migration [24]. Later, genetic and cytological analysis have further implicated astral microtubules and the cell cycle oscillator in the control of cortical migration [3]. Nuclear movements are sensitive to the level of Cyclin B-Cdk1, as revealed by genetic manipulations that alter the copy number of cyclin B [52]. Staining of microtubules in fixed embryos during cortical migration led to the idea that aster-aster repulsion drives this process. Recent live imaging experiments using embryo extracts have strengthened the evidence for this mechanism [14]. This is also supported by the observations that, in embryos in which the activity of molecular motors (Klp61F/Kinesin-5 and Klp3A) or a microtubule cross linker (Feo/Prc1) is knocked down, the process of cortical migration is significantly impaired [18]. However, it is likely that these proteins also play a role in the mitotic spindle. The ability of centrosomes to potentially organize cortical migration was supported by experiments arresting the cell cycle pharmacologically or genetically [47, 25]. In these embryos, while nuclei are arrested in interphase, centrosomes continue to replicate and separate, and eventually a fraction of them migrates to the cortex [47, 25], arguing that astral microtubules have an intrinsic ability to space centrosomes and drive their migration. However, it is unclear whether the mechanisms of this localization are what drives the process of cortical migration in normal conditions, as the significant lengthening of interphase might impact microtubule dynamics in multiple ways.

Collectively, previous observations point to the need of visualizing and quantifying nuclear migration in intact, living embryos. The major limitation towards this goal has been the difficulty of visualizing the entire nuclear population when nuclei are deep inside the embryo. Moreover, since cortical migration proceeds rapidly, this three-dimensional (3D) problem requires microscopy with high acquisition speed to cover the large size of the embryo (*∼* 0.5 mm along the Anterior-Posterior axis and 0.2 mm along the two orthogonal axes). To overcome these problems, we used light-sheet microscopy to investigate the behaviour of nuclei during cortical migration. We found that the embryo-yolk fate decision is controlled by spindle orientation, which in turn is governed by topological interactions among asters/spindles. Using ideas similar to those originally developed to describe the behaviour of foams [60], we develop a general theory for how topological interactions can become dominant in cytoskeletal systems and use genetic manipulations of the cell cycle to both confirm our theory and gain insight into the importance of this process for proper control of the nuclear density.

## RESULTS

### Visualization and quantification of nuclear movements reveal a collective and bimodal mode of migration

To capture the 3D dynamics of the nuclei migrating from the bulk to the cortex, we used light sheet microscopy to image living *Drosophila* embryos that express a fluorescently tagged nuclear pore protein, mRFP-Nup107 [33]. Combined with multi-view light sheet microscopy [35], this fluorescent construct allowed us to visualize and reconstruct the movements of almost all the nuclei in the embryo during both interphase and mitosis. Using the machine-learning based software ilastik and our own computational tools (see Methods), we segmented and tracked individual nuclei during the migrating stages over several nuclear cycles (Video S1 and Fig.S1A). Consistent with previous observations [3, 24], we found that most nuclei moved towards the cortex in a highly synchronized stepwise manner coupled to the cell cycle (Fig. 1B and 1C). Each step of migration took place over about 3 minutes, starting at the onset of anaphase and culminating in early interphase, which is longer than previously inferred from fixed embryos [24]. The fastest movement occurred during anaphase/telophase at the time of chromosome separation (Fig.1C, 1D vertical line), but residual cortex-directed movements persisted even after chromosome separation and nuclear envelope formation, as nuclei continued to position themselves after division (Fig.1C). Segmentation and tracking of individual nuclei revealed that they organize in an expanding shell during migration (Video S2). By computationally detecting the nuclei sitting on the outermost layer of such shells (Fig.1C) and projecting their velocity onto the cortical direction (Fig.S1B), we found that most nuclei move towards the cortex in a uniform manner (Fig.1D and S1C-D), while a fraction of them move toward the center of the embryo (Fig.1C, red curves, Fig.S1E). Most nuclei (more than than 90%) were sitting on the outermost shell at cycle 8, whereas this number went down progressively over the migrating phase, reaching about 70% at cycle 10 (Fig.2A). This observation indicates that some nuclei do not migrate with the outer shell but remain inside the embryo [24], where they become yolk nuclei. We thus sought to establish whether the differentiation of yolk nuclei can be closely linked to their early positioning by analyzing available single nuclei sequencing data [1], acquired from precisely staged embryos at the onset of MZT (early to mid cycle 14). Clustering analysis showed that yolk nuclei have already differentiated at the onset of MZT (Fig.1E-F, mapped to separate fate cluster), which could be identified by the marker *sisterless A* (*sisA*). Analysis of the top differentially expressed genes confirmed the upregulation of *sisterless A* (*sisA*), *spooky* (*spo*) and *serpent* (*srp*) in yolk nuclei, see Fig.S1F. These observations suggest that yolk nuclei are specified by their positioning in early embryogenesis and that such positioning drives the expression of many important developmental genes. Thus, we set out to investigate what determines which nuclei migrate to the cortex and which ones remain inside. Since both modes of migration coincide with mitotic exit, we envisioned that they might be controlled by the orientation of the mitotic spindle during nuclear divisions.

**Fig. 2.**
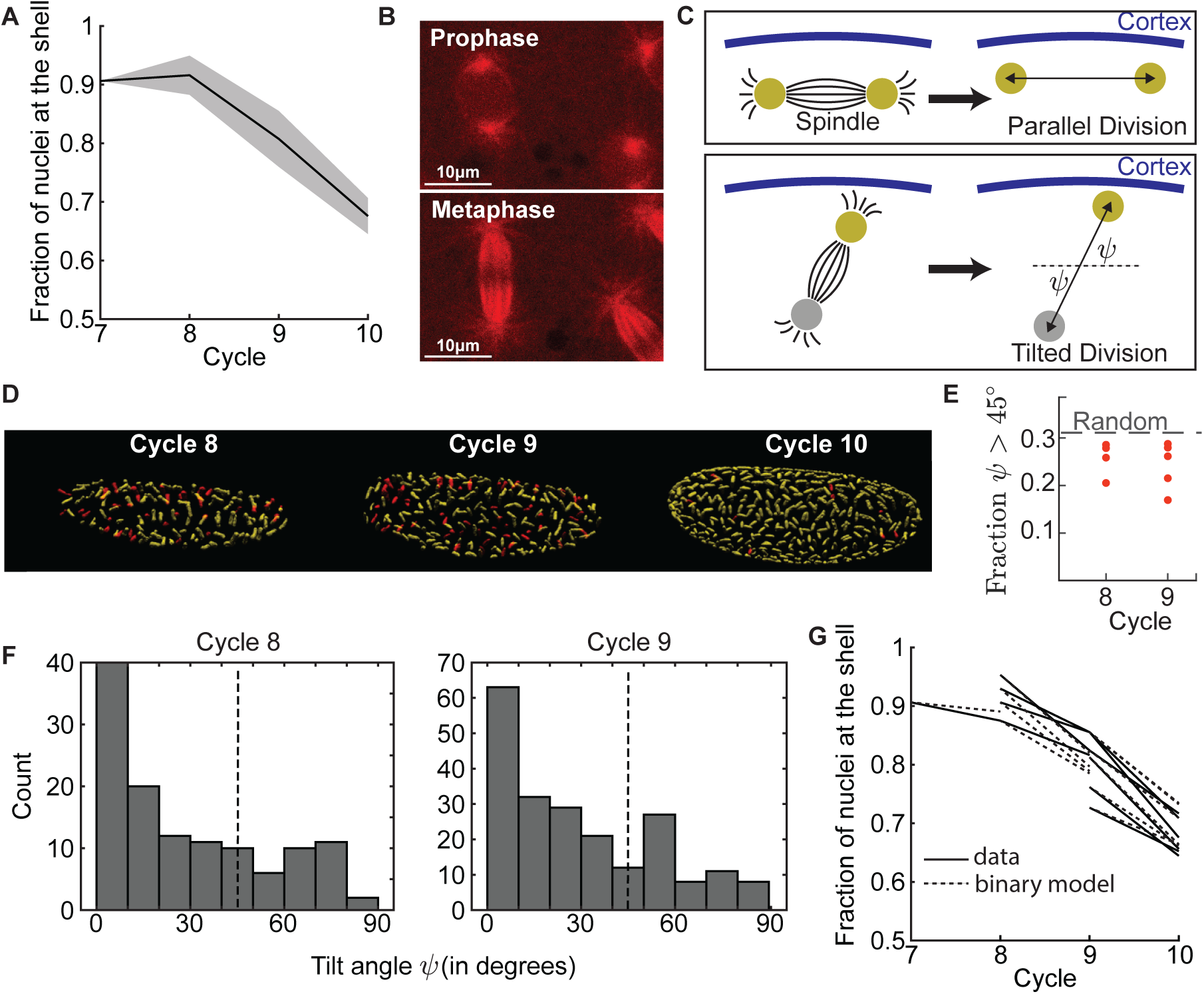
Spindle orientation drives nuclear positioning. **A,** Fraction of nuclei found at the outer nuclear shell with respect to the total number of nuclei (2*^n−^*^1^ for the *n*^th^ cycle) is shown for different cell cycle phases. **B,** Confocal image of an embryo expressing tau-mCherry, where microtubule asters(prophase) and spindle (metaphase) are visible. **C,** Motion of daughter nuclei is shown for different orientations of spindle with respect to the embryo cortex. The tilt angle *ψ* of the spindle with respect to the cortex is indicated. Parallel spindle with *ψ <* 45° leads to movement of both daughter nuclei towards the cortex, while for perpendicular spindles (*ψ >* 45°) only one of the daughter nuclei move towards the cortex. **D,** Segmented spindles (yellow and red) are shown for cell cycle 8-10. Spindle with *ψ >* 45° are marked red. **E,** Fraction of perpendicular spindles with *ψ >* 45° is shown for cycle 8 and 9. The dashed line indicates the random value 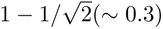 obtained for random orientations of the spindles. **F,** Frequency distribution of tilt angle *ψ* is shown for an exemplary embryo, for cell cycle 8,9 (See Fig.S2A for cycle 10). **G,** Fraction of nuclei found at the outer shell for individual embryos (black solid line) is shown for different cell cycles, in comparison to expected number of nuclei (dashed line, binary model) based on their tilt angle (= 2×fraction of parallel spindles+ fraction of perpendicular spindles).

### Spindle orientation controls nuclear fate

To capture microtubule dynamics during cortical migration, we generated a new transgenic fly line which expresses the tau microtubule binding domain tagged with mCherry (hereafter TMBD-mCherry, adapted from [23, 44]) from a maternal tubulin promoter. This fluorescent construct decorated both the spindle and astral microtubules very brightly (Fig.2B), allowing us to accurately measure the orientation of the mitotic spindle from light sheet microscopy. For each cycle, we segmented the mitotic spindles at chromosome separation (anaphase onset) and quantified their tilt angle *ψ* with respect to the embryo cortex (Fig. 2C), where *ψ* = 0° corresponds to a tangential and *ψ* = 90° to a perpendicular configuration. On any surface such as the nuclear shell described above, there are three orthogonal axes/degrees of freedom, two tangential to the surface and one normal. By random chance, the likelihood that a spindle orients more towards the normal axis, hence *ψ >* 45°, is 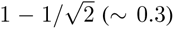, which is the probability that two random directions (the spindle and the normal axis) in 3D form an angle *>* 45° (dashed line, Fig.2E). The distribution of tilt angles (Video S3, Fig.2D) showed that the fraction of perpendicular spindles (Fig.2E) were smaller than the random value, revealing a statistical bias for tangential spindles (yellow) in cycle 8-9.

Next, we investigated how spindle orientation impacts nuclear fate determination and found that for tangential spindles, both daughter nuclei move towards the cortex and retain their position in the shell which eventually contributes to the blastoderm. On the contrary, for perpendicular spindles one daughter nucleus migrates with the outer shell while the other moves towards the bulk and becomes a yolk nucleus (Fig.2F,2G). Thus, the first fate decision in the embryo is driven primarily by spindle orientation, which determines the physical location of the nuclei.

In addition to its importance in specifying nuclear fate, proper control of spindle orientation is required to maintain the nuclear shell-like structure that drives cortical migration. Tangential spindles comprising the majority of the population ensure that nuclei expand in a shell-like architecture, since for a random distribution the movement of daughter nuclei would be incoherent, thus leading to a nuclear cloud rather than a shell. Once the nuclei reach the embryo cortex at cycle 10, spindles are found to be tangential, indicating an anchoring effect of the cortex (Fig.S2A) [6, 55]. Thus, our experiments demonstrate a central role for spindle orientation in the control of the number of yolk nuclei and nuclei that form the blastoderm, which eventually gives rise to embryonic structures.

To understand how the spindle orientation is set, we used our imaging data to detect and segment centrosomes and associated asters starting from the onset of spindle formation to chromosome separation. Tracking of these structures over a 2-3-minute window, showed negligible change of the orientation over time (Fig.S2B-C). This shows that the orientation of the spindle is not controlled by interaction among growing spindles during anaphase, but rather it is determined by centrosome orientation prior to mitosis, possibly via interaction of centrosomal asters.

### Mechanics of aster-aster interactions and inflation of the nuclear shell

To investigate how interactions among microtubule asters at different stages of the cell cycle can give rise to nuclear shell inflation, we formulated a minimal physical model assuming that asters interact via pushing forces acting on antiparallel overlaps of microtubules [12] and taking into account the geometrical organization of asters. We consider an astral unit composed of two radial microtubule asters emanating from the centrosomes attached to the nuclei [64, 56]. At proximity, asters from different astral units can interpenetrate and as a result have regions of overlapping antiparallel microtubules, with a maximum aperture of *ϕ*. The microtubule cross linker PRC1/Feo, as well as molecular motors, such as Kinesin-5/Klp61F and Klp3A, are known to act on such antiparallel microtubule pairs by sliding to generate pushing forces, which is argued to be the main driver of aster-aster interactions [54, 13]. The amount of pushing force between interacting asters depends on the amount of overlap and is set by the relative size of the asters (*l_m_*) with respect to the internuclear distance (*d*) (Fig.3A). If the asters are shorter compared to the distance between nuclei (*d/l_m_ >* 1), there is no overlap and asters do not interact. When the distance between the asters is smaller than aster length *r* = *d/l_m_ <* 1, the pushing force at the overlap region induces a separation between the nuclei. The mechanics of this separation is given by the balance of forces, which mathematically can be described as a relaxation phenomenon in a potential *W* (*r*):

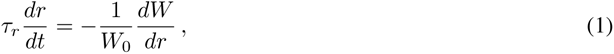

where *τ_r_* and *W*_0_ are characteristic timescale and energy scale, respectively (see Supplementary Information). The potential *W* (*r*) monotonically decays from *r* = 0 to *r* = 1 and is depicted graphically in Fig. 3B.

**Fig. 3.**
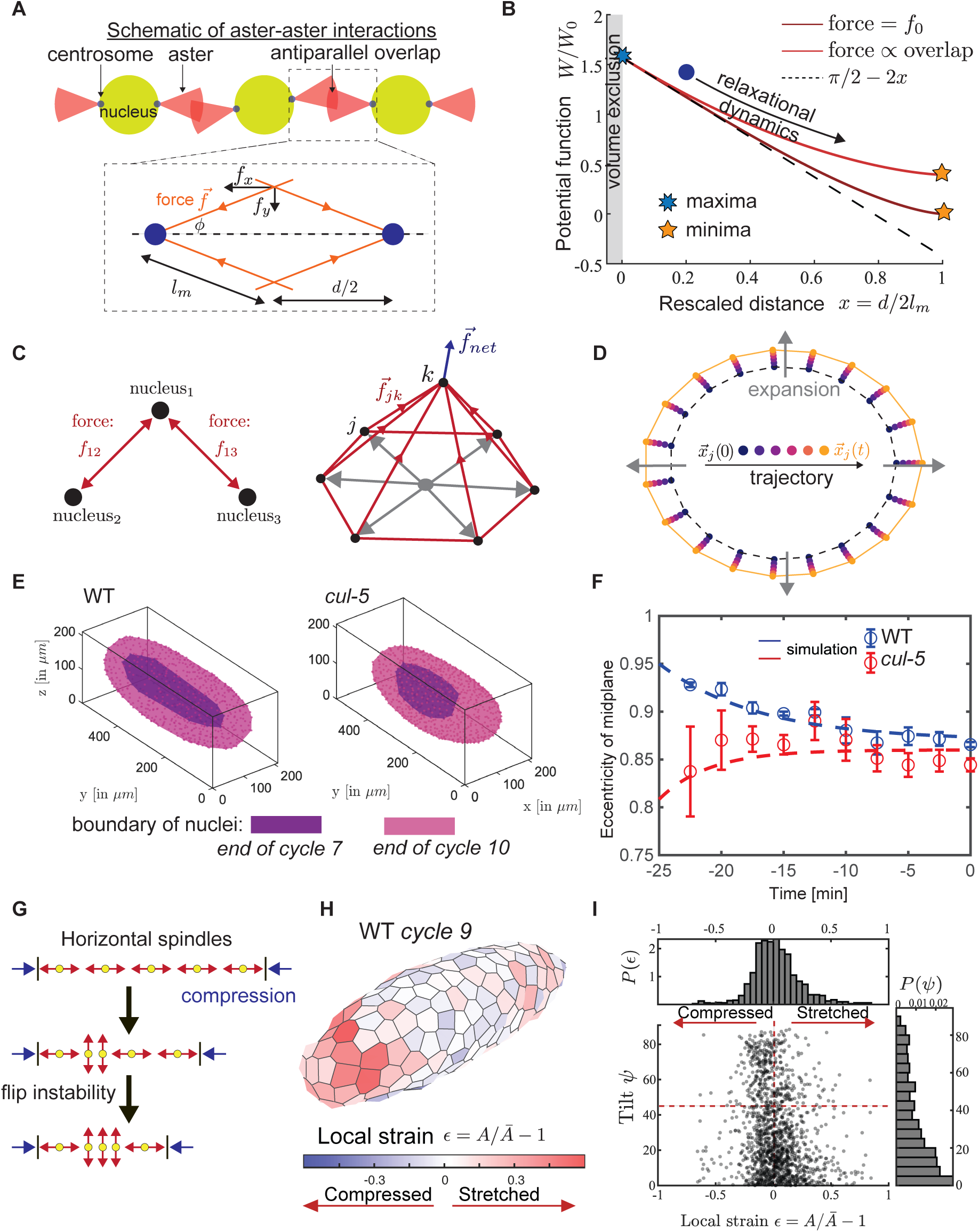
Geometry and mechanics of cortical migration via local force balance. **A,** Schematic diagram indicating the geometry of centrosomal asters and microtubule overlap regions is shown. Inset depicts the body force diagram for a microtubule pair from neighbouring asters, the pushing force *f⃗* is directed along the microtubule and can be decomposed into a pushing force along the axis *f_x_* = *f⃗* cos *ϕ* and orthogonal component *f_y_* = *f⃗* sin *ϕ*. **B,** The pushing dynamics can be as a relaxation in the potential *W* shown. The potential gradually decreases as the relative distance *x* = *d/*2*l_m_* increases between asters reaching minimum at *x* = 1, beyond which no overlap is possible. Two different cases are considered: i) where the exerted force is saturated at a constant value *f*_0_ and ii) where the force *f* is proportional to the size of the overlap. *d, l_m_* denote respectively the distance between asters and maximal length of the microtubule (shown in Fig. 3A). **C,** Force balance on a curved geometry is depicted. Pushing forces from all neighbours of *k*^th^ nucleus on a curved geometry leads to a net normal force *f⃗*_net_. **D,** The net normal force on an ellipse leads to inflation of an ellipse. Simulation results are depicted. Colorbar indicates progression. **E,** Reconstructed nuclei shells are shown from exemplary movies for cycle 7 and 10 in WT embryos and *cul-5* mutants, emphasising the difference in eccentricities in cycle 7 stage. **F,** Eccentricity of the mid-plane is plotted across time, along with fits obtained by simulations for WT and *cul-5* mutants. **G,** The mechanism of a compression induced flip instability is schematised. In this mechanism strong link between local compression and tilt is indicated. **H,** A spatial color map of relative area strain *ɛ* (see Methods) is shown on a curved Voronoi diagram for a WT embryo in cycle 9. **I,** Tilt angle *ψ* is plotted against relative strain *ɛ*. No strong correlation is found (see Fig.S4 D-E, methods) between tilt and compression.

Analysis of this mathematical model for asters distributed on a curved surface resembling the observed nuclear shells argues that pushing forces can lead to expansion of the shell, due to a net normal force (Fig.3C). This insight was confirmed with a 2D particle-based simulation where we implemented pushing forces between neighbouring astral units derived from *W* (*r*) in an ellipse. Our simulations show that the dynamics indeed leads to expansion of the shell/periphery (Fig. 3D). A simple deduction from this proposed physical mechanism is that an increase of eccentricity should follow the inflation, since higher curvature leads to higher normal force (see Supplementary Information). On an ellipsoid, the poles have higher curvature than the equatorial regions, hence leading to a larger normal net force than the tangential one. As a result, the particles at the poles move faster outward than the ones at the equator, leading to an elongation of the shell and increase in eccentricity (See Video S4). However, in wild-type embryos the expected increase of eccentricity is masked by boundary effects. Indeed, following the cytoplasmic flows at cycles 6, polar nuclei are much closer to the cortex than equatorial ones. The resulting constraints imposed by the boundary lead to an overall reduction in the eccentricity of the shell from cycle 7 to cycle 10 (see Figs. 3E-F). In our simulations, the effect of the boundary is captured by introducing a short-range potential near the boundary. In order to uncouple the effect of boundary, we turned to *cullin-5* mutant embryos. For these mutants, cytoplasmic flows are strongly reduced [28] and the nuclear shell is smaller than in WT and close to a spherical shape at cycle 7, much in contrast to the wild-type scenario. As predicted by our pushing-driven inflation model and simulations, we find that in *cul-5* mutant embryos the nuclear shell increases its eccentricity from cycle 7 to cycle 10. (Fig.3E-F and Fig.S3A,B,C) until it approaches the boundary. Thus, we conclude that experimental observations are consistent with the expansion of the nuclear shell emerging from pushing forces among asters.

### Spindle orientation is not controlled by inter-nuclear distance

We next asked whether pushing forces between neighbors in a closed shell could give rise to a mechanical instability akin to localized buckling and result in perpendicularly-oriented spindles. The resulting consequence would be that, upon compression beyond a threshold, a spindle would flip from a tangential configuration to a perpendicular one to accommodate for space (see Supplement, Fig.3G). In that case, the tilt angle *ψ* would be strongly correlated with the space available for each spindle, or local strain. To test this, we triangulated the spindles to define a Voronoi tessellation on the curved nuclear shell (see Fig.S4A-B). This allowed us to define a domain area *A* associated with each spindle as a proxy for available space. For a closed surface the total area is fixed, hence also the average area *Ā* available to each spindle/particle. We define a local strain *ɛ* = *A/Ā −* 1, where negative and positive *ɛ* corresponds to compression and dilation, respectively (see Fig. 3H). No clear correlation between tilt angle *ψ* and local strain *ɛ* emerged (see Fig.3I and Fig.S4D,E). Moreover, investigating the statistics of *ɛ*, we find that the probability distribution *P*(*ɛ*) is almost symmetric around 0 and roughly half of spindle domains are compressed, while the fraction of perpendicular spindles (*ψ >* 45°) remains below the random value bound 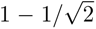 (see Fig. 2E). This indicates that distance-dependent pushing forces likely do not give rise to perpendicular spindles, on the contrary the low fraction of perpendicular spindles hints at possible alignment effects giving rise to a higher fraction of tangential spindles.

### Aster-aster alignment interactions are scale-independent and topological

Alignment interactions are ubiquitous in active biological systems that constitute anisotropic objects. These alignment interactions can emerge, for example, from contact-based or hydrodynamic effects, but in both cases the strength of the alignment decays with distance [8, 15, 41]. This raises an apparent paradox in the context of spindle alignment: since the distance between neighboring astral units increases as they push each other to drive inflation/cortical migration, the alignment interactions should progressively weaken. How could interactions still be effective and produce the observed alignment?

To resolve this paradox, we investigated the alignment effects that could emerge out of pushing forces at the antiparallel microtubule overlap regions where different asters meet. We consider a simple setup of three identical astral units (see Fig.3A), all perfectly aligned in the central plane. Activity of Kinesin-5 has been shown to create alignment in antiparallel microtubule overlaps in the context of bundling [63]. We assume that a similar mechanism operates in our biological context. If we introduce a tilt *ψ* for the central astral unit, we create an asymmetry between the amount of force being generated above and below the central plane (see Figs. 3A,4A). Assuming aligning forces at the astral overlap, for a small amount of tilt *ψ >* 0, the downward force *f_↓_* is larger than the upward force *f_↑_*, hence giving rise to an aligning/resetting torque. On the other hand, if *ψ* = 90° there are no overlaps, hence no alignment torque.

**Fig. 4.**
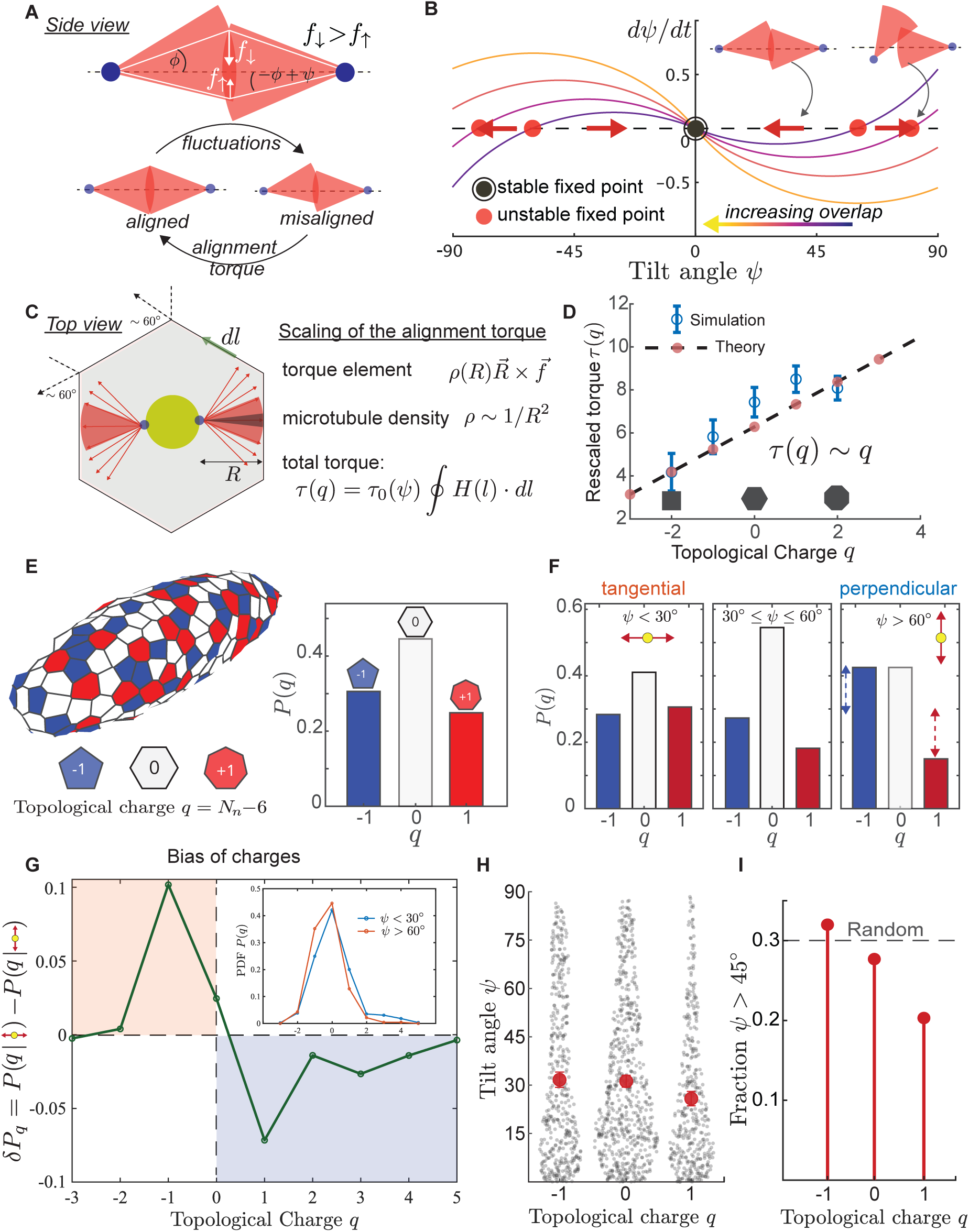
Theoretical model predicts aster-aster alignment interactions are scale-independent and topological. **A,** *Top,* Schematic force diagram is shown for neighbouring asters with a relative tilt *ψ*. Due to the asymmetry between above and below the equator,the downward force *f_↓_* is larger than *f_↑_*. Hence a net torque is exerted that reduces the tilt *ψ*. *Bottom,* this mechanism can lead to a symmetry restoration and scheme is shown. **B,** A phase portrait (*ψ, dψ/dt*) is plotted for various values of compression to determine the stability. *ψ* = 0°, 90° are found to be stable fixed points (black cricle), separated by unstable fixed points (red circle). **C,** A schematic depicts the integration of torque for an astral unit across its’ neighbours. Lengsthscales and associated scaling is shown. **D,** Rescaled magnitude of torque is plotted against the topology of the polygonal domains, for analytical prediction (red-dot and dashed line) and stochastic simulations (blue circles with error bars). Rescaled torque *τ* is found to be proportional to topological charge *q* = #neighbour-6. **E,** Distribution of *q* is shown for a WT embryo during cycle 9. **F,** Distribution function *P*(*q*) is shown for three categories of tilt angles (tangential, intermediate and perpendicular). The relative frequency of negative charges (*q* = −1) is found to increase with tilt angle *ψ*. **G,** The difference in frequency of *q* (shown in inset) between perpendicular spindles *ψ >* 60° and tangential spindles *ψ <* 30°is shown. **H,** The distribution of tilt angles is shown for *q* = 0,±1. Mean tilt angle is marked with red dot. **I,** The fraction of perpendicular spindles *ψ >* 45°is plotted as a function of topological charge *q*.

To further elucidate how these alignment interactions can lead to control of spindle orientation, we derive a dynamical equation for the tilt angle: *µψ̇* = *τ*(*ψ*) = *τ_m_g*(*ψ*) (see Supplementary Information, Fig. 4B) and identify the fixed points in the (*ψ, dψ/dt*) phase plane (Fig. 4B). Here, the magnitude of the alignment torque is given by *τ_m_*. Phase-plane analysis [53] shows that *ψ* = 0° is a stable fixed point (black circle Fig. 4B) with a boundary stable point at *ψ* = 90°, separated by an unstable fixed point (red circle Fig. 3B). This provides a physical mechanism of alignment where differences in tilt between neighbouring astral units can be reduced. However, the total amount of alignment torque acting on a given aster depends on the entire region of overlap for an astral unit along its periphery in the nuclear shell. To estimate the amount of total torque, we consider the geometry of the astral units on the shell and integrate over its periphery to account for all the interaction with its neighbours (Fig. 4C). For radial asters, the density of microtubules *ρ* at distance *R* from the center *ρ*(*R*) *∼* 1*/R*^2^ [34], while the unit torque (*R⃗ × f⃗*) generated by each microtubule contact *∼ R*. Hence, along the periphery of the Voronoi domain for a small arc *dl* (see Fig.4C), we can define the torque element and total torque as

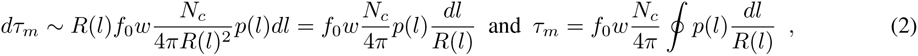

where *f*_0_ is a typical force exerted at a microtubule contact, *w* is the width of the nuclear shell and *N_c_* is the number long microtubules emanating from the centrosome. The parameter *p*(*l*) is the probability of antiparallel overlap at the location *l* around the periphery. The geometric scaling reveals that the torque is scale invariant. In other words, if we consider a nuclear shell where the thickness *w* remains the same but all the Voronoi domains (and the nuclei-to-nuclei distance as a result) are rescaled by a numerical factor *λ*, the torque *dτ_m_ ∼ d*(*λl*)*/λR*(*l*) = *dl/R*(*l*) is independent of the rescaling factor *λ*. This theoretical analysis reveals two key insights: first, that the alignment is independent of the internuclear distance in line with our findings; second, that decoupling of the size of the domains imply a dependence on the geometry/shape of the astral domain. Note that this geometric mechanism implies that the amount of alignment stays unchanged as relative distances change in the course of the inflation process, which likely contributes to the robustness of the process.

To investigate the effect of geometry, we make use of curved Voronoi tessellations and define the shape of the astral domains with an iso-perimetric coefficient 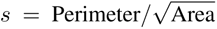 and local topological charge *q* = #neighbors *−* 6. We find that, despite spindles being rod-like nematic objects, the domains are largely isotropic with *s* = 3.8 *±* 0.1, similar to the case of a regular isotropic hexagon (see Fig.S4C, Supplementary Information). This lack of nematicity in spite of the rod-like structure of spindles argues strongly that during cortical migration the nuclei/spindles do not experience a very compressed regime, as that would have driven the spindles to acquire nematic organization leading to alignment. Next, we performed stochastic simulations to estimate the integral of the alignment torque around the periphery of domains drawn from the experimental data. We consider the orientation and the up-down degree of freedom for the domain and its neighbours (see Methods for details). From the simulations we find that the estimated torque scales with the topological charge *q* and does not depend on the size of the domains (see Fig. 4D). To provide an intuition for this result, let us consider a regular polygon with *n* sides where the asters fan out with an aperture of *∼* 120°. The integral of the torque element *p*(*l*)*dl/R*(*l*) is a number *O*(1), which counts how many sides of the polygon interact with the astral unit. The average number of sides that fall within the two astral fans of aperture *α* is proportional to the number of neighbours *n* (see Supplement). For each interacting side, the corresponding neighbouring aster can point in the correct configuration with probability *p*. Hence the amount of torque *τ_m_* is proportional to the number of neighbours *n* and thereby to the topological charge *q*. An alternative way to derive this relationship is to consider the mean-field case *p*(*l*) = *p* and make use of a well-known identity in integral geometry and mechanics of foams and soap films [50, 40]:

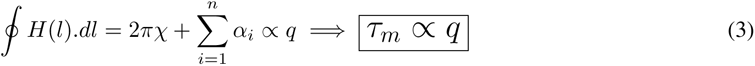

where *H* = 1*/R* is the curvature, *χ* is the Euler characteristic and *α* is the exterior angle as indicated in Fig.3C.

Eq.(3) is the 2D version of the Gauss-Bonnet theorem that shows that integral curvature 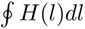 is a topological invariant (see Supplement for discussion) applied to polygons [17]. This identity was also previously used by von Neumann to derive the eponymously-named topological growth law of a 2D foam cell [60]. Similar to von Neumann’s law, we reach the central conclusion that the magnitude of alignment torque of an astral domain is independent of its size but is linearly proportional to the topology of the neighbourhood (see Fig.4D), indicating that higher number of neighbours imply more alignment.

To test the predicted relationship between spindle orientation and topology, we investigated the distribution of topological charges *q* in WT embryos at cycle 9 (n=8, see Fig.4E). For simplicity and reliable statistics, we here group all positive charges (*q >* 0 or more than 6 neighbors) as +1 and negatives (*q <* 0 or less than 6 neighbors) as *−*1. We find that perpendicular spindles are more likely to be negatively charged (*q* = *−*1) than tangential spindles (see Fig.4F,4G), indicating a correlation between higher tilt angle *ψ* and negative charge *q*. As a complement, we also investigated the distribution of tilt angles *ψ* as a function of charge *q*. While the average tilt angles *ψ̄* (indicated by red circle with errorbars in Fig.4H) differ only mildly for different *q*, the fraction of spindles with *ψ >* 45°depends strongly on *q* (see Fig. 4I). For *q* = *−*1, the fraction of perpendicular spindles is the highest (close to the random value) and decreases as *q* increases. Together, these results establish a strong bijective relation between spindle orientation and topological charge, as predicted by our theory. Furthermore, our analysis provides general theoretical insights on how topology emerges as the dominant regulator of spindle orientation.

### Cell cycle mutants establish scale-independent and topological nature of alignment interactions

To further test the scale-independent and topological nature of alignment interactions in determining spindle orientation, we sought to tune the inter-nuclear distances with the aid of genetic manipulations of the cell cycle. The two main processes that set the average inter-nuclear distance are: 1) the cell cycle dependent pushing forces, which drive the inflation and, as a result, the increase of the distance among nuclei; 2) the nuclear divisions, which increase the density and decrease the average distance. Changing the duration of the cell cycle, and more specifically the duration of the period of nuclear migration, with various mutants gives us a way to tune inter-nuclear distances. The stepwise migration of the nuclei takes place from anaphase to early interphase (Fig. 1C). Previous literature indicates that cortical migration is sensitive to Cyclin-dependent kinase 1 (Cdk1) activity [3], which our analysis of nuclear speed and emission ratio of Cdk1/PP1 FRET sensor confirms (Fig.5A). Specifically, it was shown that Cdk1 activity limits cortical migration and it was suggested that this is due to Cdk1 inhibition of astral microtubule growth [52, 58]. To manipulate Cdk1 activity, we generated flies that have different copy numbers of *cyclin B* (*cycB*) (see Methods), which is a dominant binding partner of Cdk1 in the early embryo and can control Cdk1 activity in a dosage-sensitive manner [11, 31, 32]. Live imaging showed that, as previously reported from fixed embryos, the nuclei in *1x cycB* embryos reach the cortex early, whereas *6x cycB* embryos showed delayed migration (Fig.5D). In addition, we screened heterozygous cell cycle mutants for their nuclear migration defects and found that in *polo* (PLK1) heterozygous embryos (hereafter *polo/+*), the nuclei also reached the cortex earlier than in WT (Fig.5D). We also found that, while there were no discernible differences between different mutants in terms of the number of nuclei at the shell at cycle 9, stark differences in nuclei number emerge by cycle 11 (see Fig.5B,C), due to the earlier cortical anchoring of nuclei that reach the cortex at cycle 9 instead of cycle 10.This leads to different nuclear densities in mutants as well as nuclear-to-cytoplasmic ratio, which is a known modulator of the cell cycle prior to the maternal-to-zygotic transition [38].

**Fig. 5.**
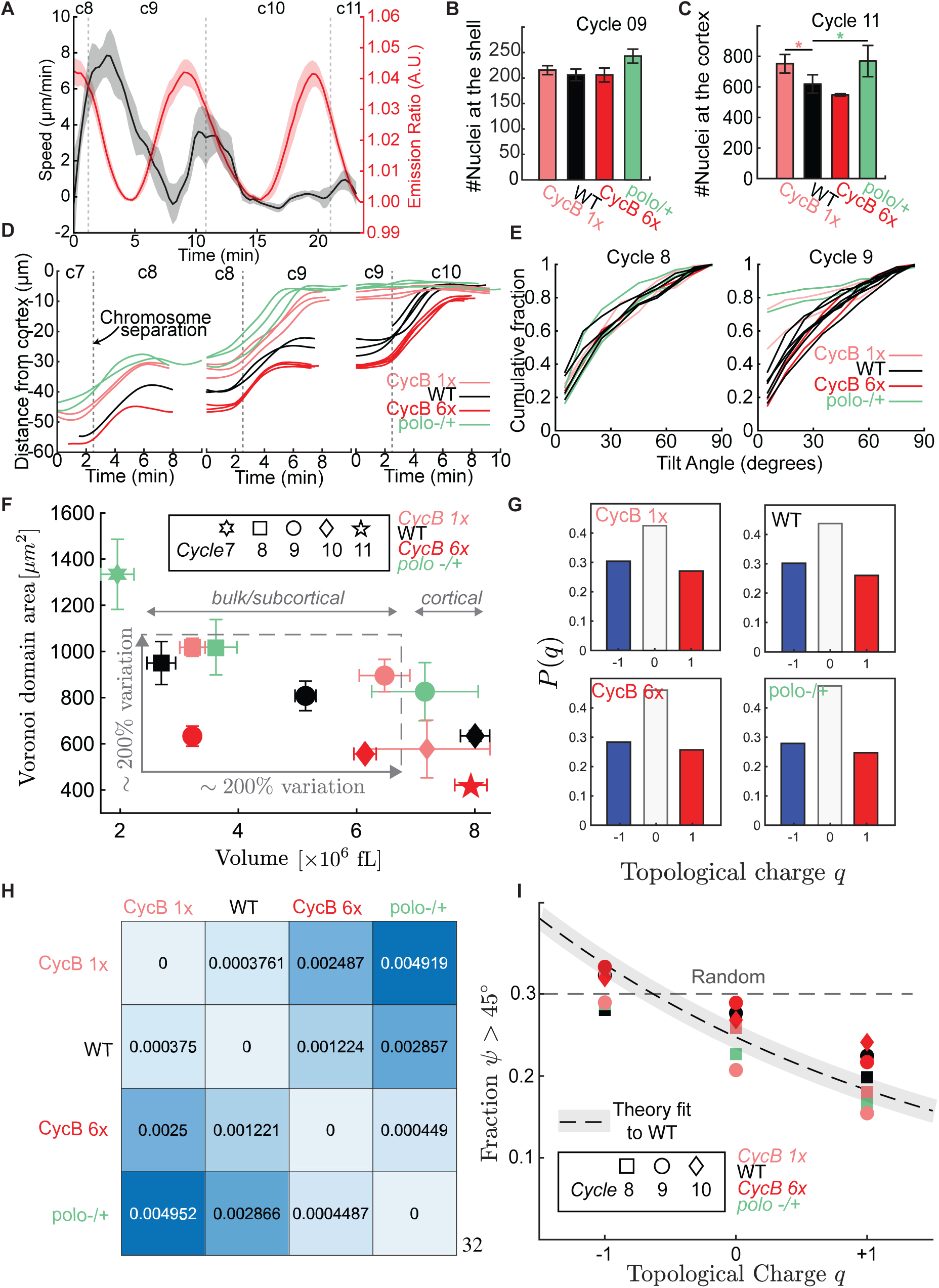
Modulation of internuclear distances using cell cycle manipulations confirms scale-independent and robust aster-aster alignment. **A,** Nuclear speed (black line=mean and shaded region= standard deviation) and emission ratio of Cdk1/PP1 FRET sensor is shown during cell cycle 08-11. **B-C,** The number of nuclei at the outer shell is shown for WT and various cell cycle mutants (errorbars indicate standard deviation) during cell cycle 09 and 11. **D,** Average distance between nuclei and the embryo cortex is shown across cell cycles, for WT and cell cycle mutants. The cell cycle mutants exhibit various duration of cell cycle and associated degree of inflation. **E,** Cumulative distribution function of tilt angles is plotted for WT and cell cycle mutants in cycle 8 and cycle 9. **F,** A plot is shown for average Voronoi domain area associated with astral units and volume of the outershell is plotted for WT and cell cycle mutants across different cell cycles. A cortical and bulk region is marked. **G,** Distribution of charges *P*(*q*) is shown for cycle 09 WT, Cyc1xB and Cyc6xB mutant embryos as well as cycle 08 heteozygous polo-/+. No marked difference is found. **H,** Relative entropy/Kullback-Leibler divergence is calculated and shown for distributions in panel **G**. Relative entropies are found to be negligible ~ 0. **I,** Fraction of perpendicular spindles (*ψ >* 45°) is plotted against topological charge *q* for WT and mutant embryos with varying domain sizes (see panel **F**) and cell cycle stages. For comparison theoretical fit is shown (black dashed line with shaded region).

As for the tilt angle *ψ*, we found that its cumulative distribution was only different in cases where the nuclei have already reached (and are anchored to) the cortex (see Fig.5D,E). Thus, we focused on datasets for WT and mutant embryos where the nuclei have not yet reached the cortex and computed both the average domain area and total shell area. This dataset (indicated as bulk/subcortical in Fig.5E), contained different nuclear cycles across all the mutants and spanned a large range in terms of domain area/size (500 *−* 1000*µm*^2^, see Fig.5F). We found that despite these differences in size, the distribution of topological charges *P*(*q*) for different mutants at cycle 8 looked identical (Fig.5G) and we found no discernible difference in terms of the Kullback-Leibler divergence (Fig.5H). Most notably, we discovered that the relationship between the fraction of perpendicular spindles and topological charge holds true across the dataset from different mutants with varying domain area and progression of cortical migration and agrees with theoretical predictions (Fig.5I). In sum, the modulation of inter-nuclear distances by cell cycle manipulations establishes that spindle orientation is not dependent on size/inter-nuclear distance, but rather is set by the topology of the neighbourhood that governs the alignment interactions with the neighbouring astral units.

## Discussion

Proper control of morphogenesis requires that cellular and subcellular processes are robust to several sources of noise [61]. Here, we investigated cortical migration in the early *Drosophila* embryo to elucidate how a precise morphogenetic process arises from the integration of the cell cycle oscillator, cytoskeletal dynamics and the physical/geometrical properties of the embryos. We found that cortical migration and the embryo-yolk cell fate decision are governed by local interactions of microtubule asters/spindles that set spindle orientation at a global level. The robustness of this process is ensured by the scale-independent and topological nature of the interactions, which are insensitive to variations in the relative distance between nuclei and their dynamic changes in size and shape in the course of the cortical migration process. The proposed mechanism provides a reliable way to ensure a high fraction of tangential spindles, which is essential to retain the observed nuclear shell-like geometry during the migration process. The expanding shell-like geometry also provides a clear separation between the embryonic surface nuclei (2D) and the yolk nuclei. Such separation can further help the coordination in morphogenesis. In fact, in embryos of insects where nuclei retain a three-dimensional organization, large heterogeneities in compartmentalization as well as decoherence in cell cycle dynamics are observed [20]. Such decoherence can lead to varying nuclear domains in early embryo, much in contrast to what is seen in *Drosophila melanogaster*. In view of *Drosophila’s* need to avoid predation [46], it is likely that speed constitutes the main evolutionary drive for these features of cortical migration. We propose that the uncovered feedback mechanisms not only determine the embryo-yolk fate decision, but also provide the template for collective coordination at later stages of development.

The importance of topological interactions in cortical migration extends our understanding of how nuclear positioning is precisely controlled in *Drosophila* embryos. In previous studies, we showed how a feedback between the cell cycle oscillator and cortical actomyosin can ensure robust spreading of nuclei in cycles 4-6 via regulation of cytoplasmic flows [16, 37]. Here, we found a new mechanisms contributing to robustness based on geometric rules rather than mechano-chemical feedback. Thus, investigating the control of early embryogenesis reveals novel and surprising aspects of biological robustness. Accurate control of cellular/nuclear positioning is essential for the first fate decision in embryos of multiple species. In mouse embryos, for example, the first cell fate decision of whether a cell becomes trophectoderm or inner cell mass is linked to cell positioning [42]. Both cortical actomyosin contractility and spindle dynamics have been implicated in this process [36]. The property we have uncovered here is that the decision of whether nuclei in the *Drosophila* embryo become yolk or blastoderm nuclei is specified by spindle orientation, which in turn is controlled by topological interactions among spindles/microtubule asters. Notably, yolk nuclei are programmed to have a different gene expression program at the onset of the maternal-to-zygotic transition when they express several factors that have been implicated in developmental processes [62, 57, 45]. Thus, our work demonstrates a close link between the first fate specification in the *Drosophila* embryo and collective cytoskeletal dynamics.

Despite its crucial role for early *Drosophila* embryogenesis, the process of cortical migration had remained poorly understood. A major limitation in the study of this process was the inability to visualize nuclear movements in intact living embryos. Previous efforts to study cortical migration had relied on analysis of fixed embryos and/or on the investigations of the effects of genetic mutations on the process [3, 24, 18]. These experiments implicated the role of spindle and mitotic separation movements and aster-aster interactions in the process. The importance of the aster-aster interactions was supported by experiments arresting the cell cycle pharmacologically or genetically and demonstrating that centrosomes continued to replicate and that a subset of them migrate to the cortex [13]. Moreover, nuclear positioning is impaired in embryos with reduced activity of cross-linking proteins and motors (Feo/Prc1, Klp3A and Klp61F/Kinesin-5) likely to act on the overlapping microtubules between asters [18]. Here using a combination of light sheet microscopy and novel computational approaches, we map out the underlying physical mechanisms that govern cortical migration in 3D and give rise to the embryo-yolk nuclear fate decision. Using a new transgenic line to visualize microtubules, we were able to visualize spindles and compute their orientation during the migration process. State-of-the-art experimental data allowed us to develop in parallel a physical theory. In particular, by using integral geometry, we were able to relate spindle orientation to fundamental geometric parameters that capture the nature of interaction with neighbouring spindles/asters and show that spindle orientation is not dependent on distance to the neighboring spindles but rather on the topology of the neighborhood, that is the number of neighbors. This insight was confirmed by genetic manipulations of the cell cycle oscillator. These manipulations altered nuclear spacing without changing topology and we observed similar distributions and dependencies of spindle orientation. Notably, these experiments also demonstrated the robustness of the process of cortical migration to fluctuations in inter-nuclear distance. Thus, our work illustrates how the combination of advanced live imaging techniques, computational image analysis and theory can shed light on classic developmental questions.

A fundamental goal of theory in biology is to uncover emergent properties at a macroscopic scale from microscopic interactions and the biological function of these emergent properties. Respective advantages of metric and topological interactions have been previously discussed in the context of starling flocks [4]. Here, we have focused on development, and provided an intuitive and predictive physical framework for the emergence of topological interactions [30] from microscopic building blocks of the cytoskeleton, viz., microtubules and motors. We are positive that that ideas and methods presented here will be broadly applicable to a variety of self-organizing complex systems due to the widespread role of cytoskeletal elements in living and active matter.

## Acknowledgements

We acknowledge Stefan Gunther and Lars Hufnagel for discussion and for sharing their original observations of cortical migration. We thank the Bloomington Drosophila Stock Center, the Kyoto Drosophila Stock Center for providing stocks. We thank the Drosophila Genomics Resource Center and Jesse Gatlin for constructs. We thank Christine Field and Tim Mitchison for discussions.

## Author contributions

Conceptualization: WH, AM, SJS, MV, SD; Methodology: WH, AM, SJS; Software: WH, AM, SJS; Validation: WH, AM; Formal Analysis: WH, AM; Investigation: WH, AM, LH, ZL, AC, NPM; Resources: WH, AM, AC; Data Curation: WH, AM, ZL; Writing-Original Draft: WH, AM, SD; Writing-Review & Editing: All Authors; Visualization: WH, AM, ZL; Supervision: SJS, MV, SD; Project Administration: SJS, MV, SD; Funding Acquisition: AM, MV, SD.

## Data and materials availability

All materials are available upon request.

