## Supplementary Figures for "Scale-independent topological interactions drive the first fate decision in the *Drosophila* embryo"

**Figure. S1**

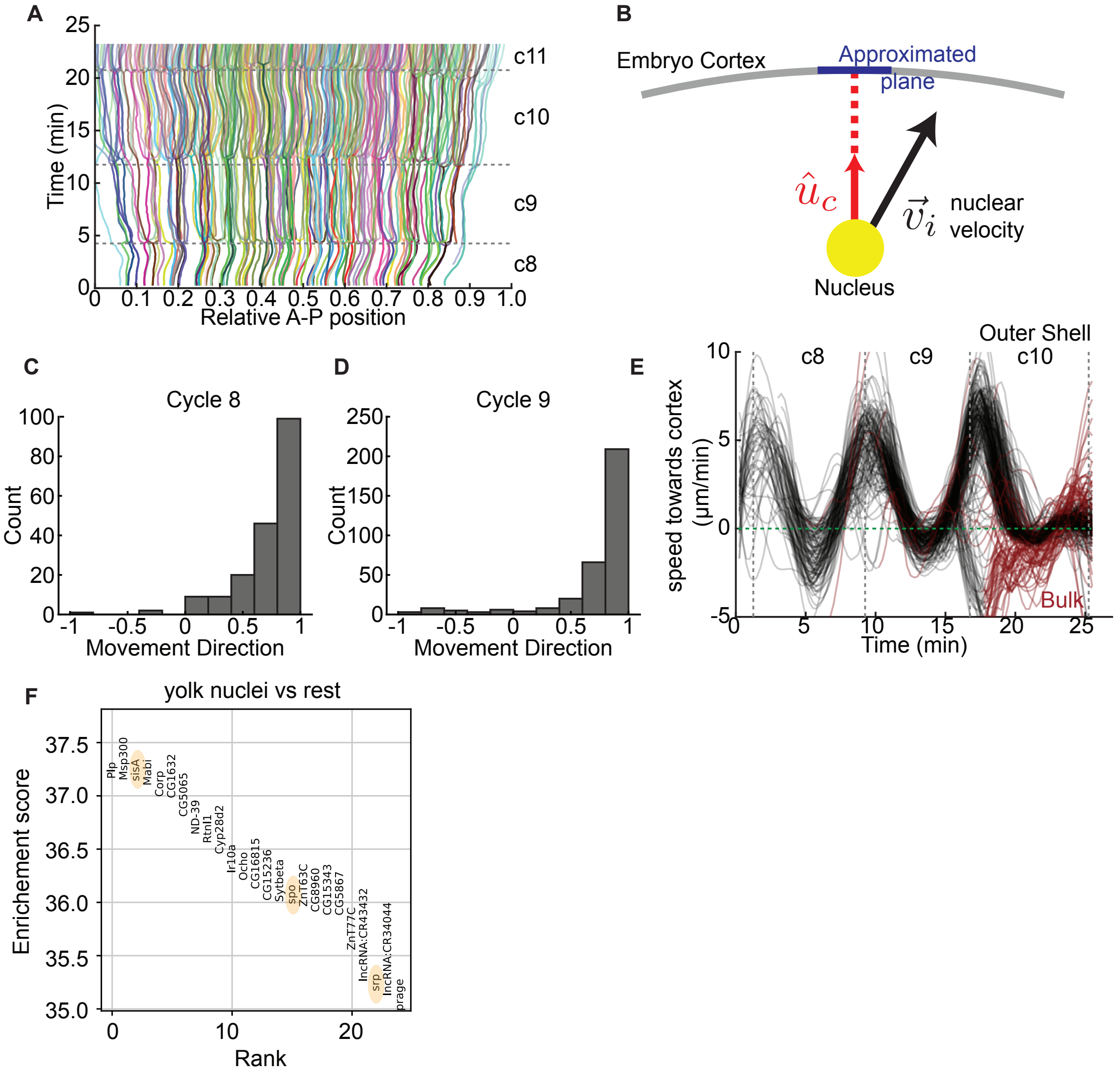
A
Time (min)
Relative A-P position
c11
c10
c9
c8
B
Embryo Cortex
Approximated plane
$\hat{u}_c$
$\vec{v}_i$
nuclear velocity
Nucleus
C
Cycle 8
Count
Movement Direction
D
Cycle 9
Count
Movement Direction
E
Outer Shell
c8
c9
c10
speed towards cortex (μm/min)
Bulk
Time (min)
F
yolk nuclei vs rest
Enrichement score
Rank
Plp
Msp300
sisA
Mabi
Corp
CG1632
CG5065
ND-39
Rtnl1
Cyp28d2
Ir10a
Ocho
CG16815
CG15236
Sytbeta
spo
ZnT63C
CG8960
CG15343
CG5867
ZnT77C
lncRNA:CR43432
srp
lncRNA:CR34044
prage

**Figure S1 A,** Lineage tracing of segmented and tracked nuclei are shown along their relative position in the anterior posterior axis from cycle 07 to cycle 11. **B,** A schematic depiction of a nuclei with respect to the embryo cortex is shown. The nuclear velocity $\vec{v}_i$, the unit orientation vector pointing to the closest plane in the cortex $\hat{u}_c$ is marked. **C,D** A frequency distribution of movement direction of the nuclei during cycle 8 and cycle 9 is shown. The movement direction is defined as the dot product of $\vec{v}_i/|\vec{v}_i|$ and $\hat{u}_c$. **E,** Nuclear speed towards the embryo cortex is shown as a function of time. **F,** Enrichment score for the top 25 expressed genes in the yolk nuclei (with respect to the rest of the nuclei) are shown in rank order. Genes *sisA*, *serp* and *spo* are highlighted.

**Figure. S2**

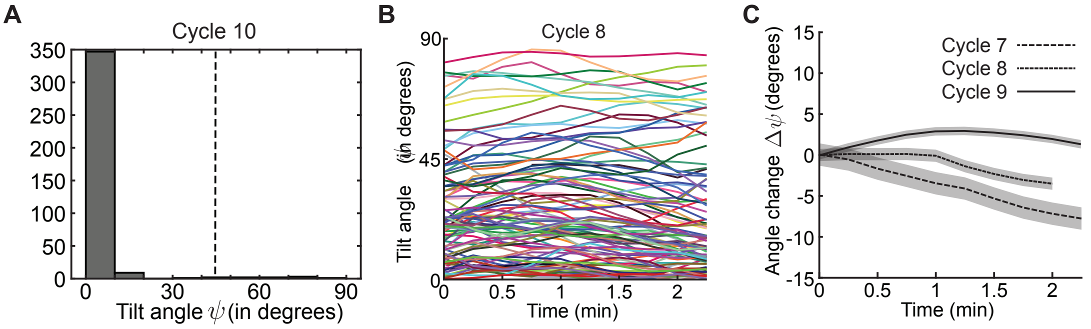
A
Cycle 10
350
300
250
200
150
100
50
0
0
30
60
90
Tilt angle ψ (in degrees)
B
Cycle 8
90
45
0
Tilt angle ψ (in degrees)
0
0.5
1
1.5
2
Time (min)
C
15
10
5
0
-5
-10
-15
Angle change Δψ (degrees)
Cycle 7
Cycle 8
Cycle 9
0
0.5
1
1.5
2
Time (min)

**Figure S2 A,** Frequency distribution of tilt angle $\psi$ is shown for an embryo during cycle 10 when they have already reached the embryo cortex. **B,** Temporal traces of tilt angle $\psi$ is shown from mid-interphase to onset of metaphase during cycle 8. **C,** The average change in tilt angle $\Delta\psi$ is shown for cycle 7,8 and 9. Shaded region marks the standard deviation.

**Figure. S3**

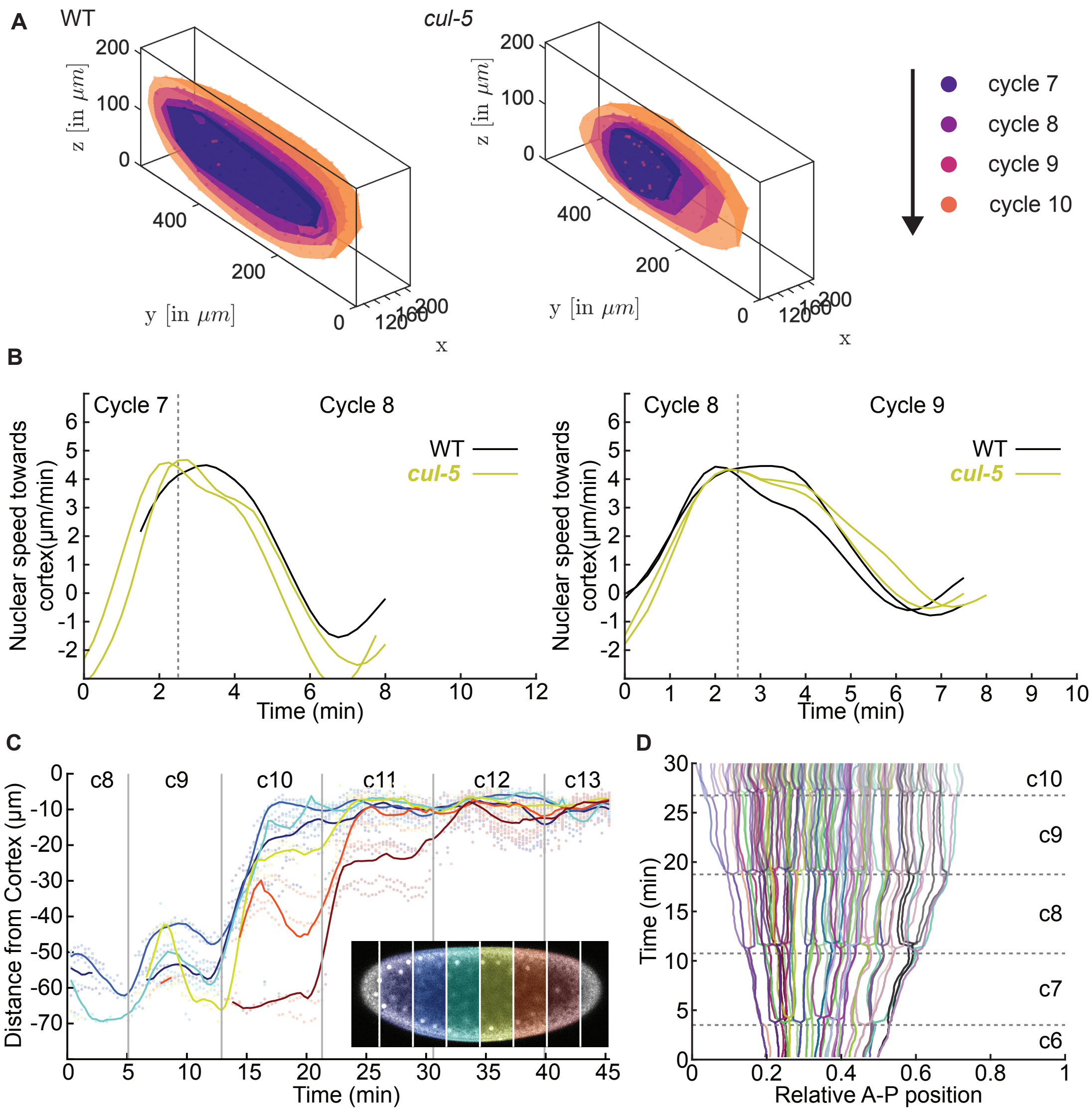
A
WT
cul-5
z [in μm]
y [in μm]
x
cycle 7
cycle 8
cycle 9
cycle 10
B
Cycle 7
Cycle 8
Cycle 9
WT
cul-5
Nuclear speed towards cortex(μm/min)
Time (min)
C
c8
c9
c10
c11
c12
c13
Distance from Cortex (μm)
Time (min)
D
c10
c9
c8
c7
c6
Time (min)
Relative A-P position

**Figure S3 A,** A hemispherical view of the nuclear shell/boundary is shown during cycle 7-10 for wildtype and *cul-5* mutants. Inflation and differences in morphology is noted. **B,** Average nuclear speed towards the embryo cortex is plotted for wildtype and *cul-5* mutants, during the transition from cycle 7 to cycle 8 (left) and from cycle 8 to cycle 9 (right). **C,** Distance of the nuclei from the cortex is shown as a function of time for a *cul-5* mutant embryo. Temporal traces (dots) and average trajectories are shown for different anterior-posterior regions (color coded). **D,** Lineage tracing of segmented and tracked nuclei are shown along their relative position in the anterior posterior axis from cycle 7 to cycle 10 for a *cul-5* mutant embryo. Note that the axial spread is smaller than a WT embryo (Fig.S1a).

**Figure S4**

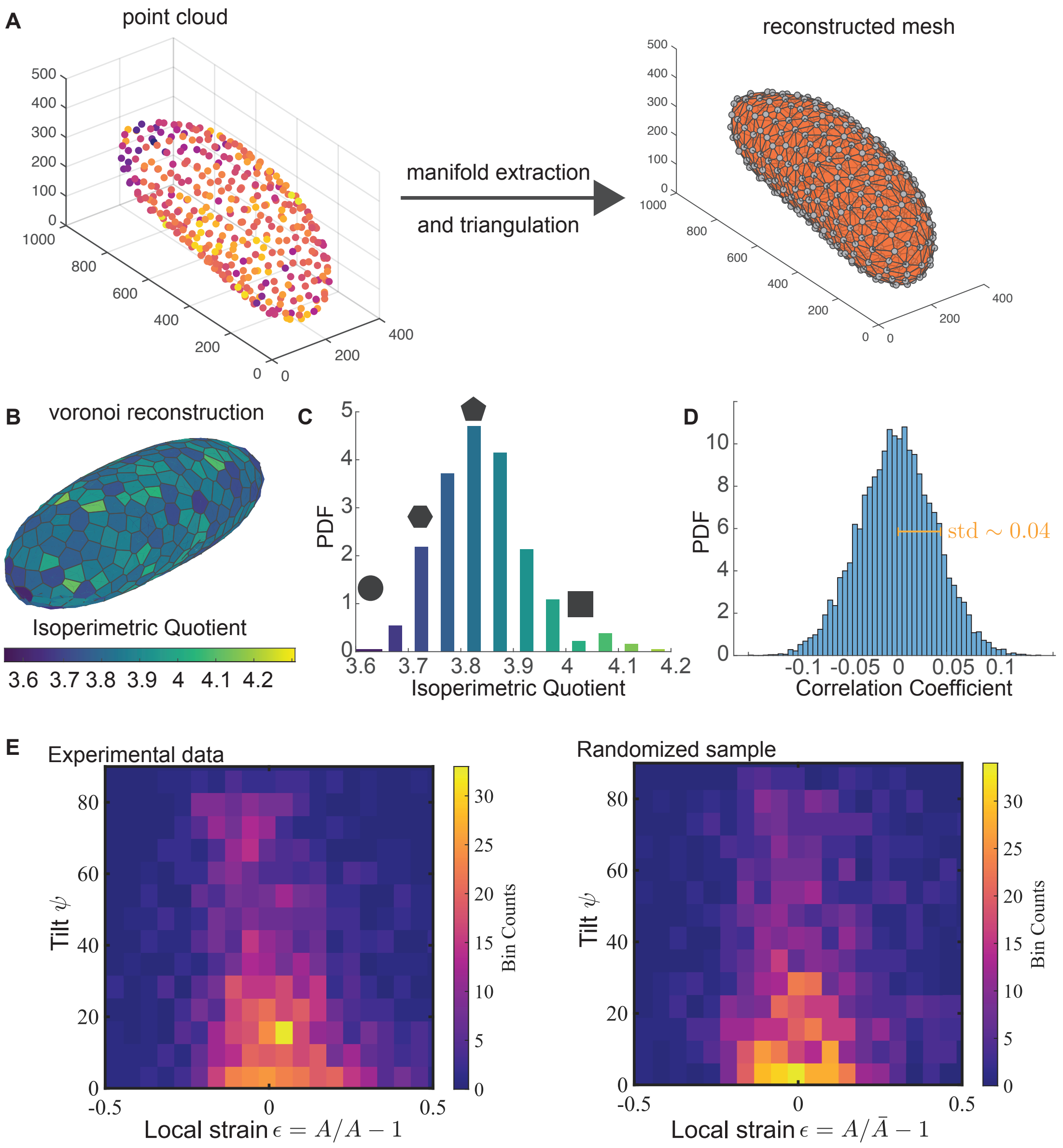
A
point cloud
500
400
300
200
100
0
1000
800
600
400
200
0
0
200
400
manifold extraction
and triangulation
reconstructed mesh
500
400
300
200
100
0
1000
800
600
400
200
0
0
200
400
B
voronoi reconstruction
Isoperimetric Quotient
3.6 3.7 3.8 3.9 4 4.1 4.2
C
PDF
5
4
3
2
1
0
3.6 3.7 3.8 3.9 4 4.1 4.2
Isoperimetric Quotient
D
PDF
10
8
6
4
2
0
std ∼ 0.04
-0.1 -0.05 0 0.05 0.1
Correlation Coefficient
E
Experimental data
Tilt ψ
80
60
40
20
0
-0.5
0
0.5
Local strain ε = A/A − 1
Bin Counts
30
25
20
15
10
5
0
Randomized sample
Tilt ψ
80
60
40
20
0
-0.5
0
0.5
Local strain ε = A/Ā − 1
Bin Counts
30
25
20
15
10
5
0

**Figure S4 A,** An example of a mesh reconstruction is shown for a nuclear point cloud data from a wildtype embryo during cycle 9. **B,** Dual/Voronoi mesh is shown for the reconstructed mesh. Facecolor indicates isoperimetric quotient (Perimeter/$\sqrt{\text{Area}}$), a measure of shape for astral domains. **C,** Probability density is shown for isoperimetric quotient for an example embryo (same as Fig.S4B), with numerically associated regular polygons/circle to emphasize an intuition of the shape. **D,** Correlation coefficient is calculated between tilt $\psi$ and negative strain $\epsilon < 0$ (see Fig.3A) after randomization. A probability distribution is plotted for the correlation coefficients upon randomization. Standard deviation of the randomization $\sigma_r$ is shown in orange. The correlation of the experimental data is found to be $\sim 2\sigma_r$. **E,** Binscatter plots for experimental data (left) of tilt angle $\psi$ and strain $\epsilon$ and a randomized sample of the experimental data (right) is shown.
