## Supplementary Information for "Scale-independent topological interactions drive the first fate decision in the *Drosophila* embryo"

#### Table of contents:

- Captions for Videos S1-S4
- Materials and Methods
- Supplementary Note

**Correspondence**
Further information and requests should be addressed to Stefano Di Talia.

**Materials Availability**

**Data and Code Availability**
The data and codes generated during this study will available openly upon publication.

#### Captions for Movies

**Video S1:**
Time lapse imaging with maximum intensity projection of an exemplary *Drosophila* embryo from cycle 07 to cycle 10 is shown with mRFP-Nup107 (red). Scale bar is $100\mu m$.

**Video S2:**
Tracing of 3D segmented nuclei (marked in variety of colours) is shown during cycle 07 to cycle 10. As the movie progresses expansion of the nuclear shell can be seen. Scale bar is $100\mu m$.

**Video S3:**
Segmented spindles are rendered and visualized in 3D. Spindles with tilt angle $< 45°$ are marked in golden , while spindles with tilt angle $\geq 45°$ are marked in red.

**Video S4:**
A typical simulation of expansion is shown where the system starts with 20 particles that divide/double and relaxes the forces by pushing on eachother. This leads to inflation as well increase in eccentricity. After relaxation, the number is again doubled and the process is repeated until interaction with the boundary.

### Materials and Methods

##### Experimental model

All experiments in this study used Drosophila embryos that are mostly in the syncytial preblastoderm or blastoderm stage (nuclear cycles 7-13). The fly lines used or generated in this study are listed in the key resources table.

##### Molecular Biology and Transgenic Flies

To generate the flies that express extra CycB, the genomic sequence of Drosophila CycB along its endogenous promoter was isolated from 10kb NheI fragment of BAC CH322-115O15. This fragment was then cloned into pCasPer4, and then injected into w1118 for random insertion transformation. We obtained 2 independent insertion lines, each on the 2nd and 3rd chromosomes. This allowed us to generate viable flies that have up to 6 copies of cyclin B, including the 2 endogenous copies (6x CycB: w; pCasPer4-CycB/pCasPer4-CycB; pCasPer4-CycB/pCasPer4-CycB). The tau microtubule binding domain (TMBD) clone was a gift from Jesse C. Gatlin, originally reported in [2] . TMBD fragment was generated from PCR, conjugated with mCherry, then cloned in the plasmid pBabr containing the maternal Tubulin promoter and the spaghetti squash 3' UTR (a gift of Yu-Chiun Wang and Eric Wieschaus, Princeton University), using Gibson assembly (NEB). The pBabr construct was then injected into C31 based fly line zh-86Fb (http://www.flyc31.org), which has the insertion site at 86Fb on the 3rd chromosome.

##### Live imaging of embryos with confocal microscopy

Flies of desired genotype were placed in a cage with an apple juice agar plate topped with a small yeast paste. After 0-2 hours, embryos were collected and dechorionated in 100% bleach for 1.5 minutes. Dechorionated embryos were then mounted on an air-permeable membrane with halocarbon oil 27 (Sigma-Aldrich, CAS Number 9002-83-9) and covered with a cover slip. Images were acquired through confocal microscope Leica SP8 and its software Leica Application Suite X (LAS X), with 600Hz scan speed, 20x magnification objective with immersion oil (HC PL APO CS2 20x/0.75 IMM), and 1.1x zoom with 800x350 pixels which gave 0.661µm/pixel resolution. No z stacks were used, and the frame rate was 1/18 sec. 458nm laser was used for Cdk1/PP1-FRET sensor, and 561nm laser was used for mRFP-Nup107 and TMBD-mCherry. HyD detectors were used for both channels.

##### Live imaging of embryos with light sheet microscopy

Flies of desired genotype were placed in a cage with an apple juice agar plate topped with a small yeast paste. After 0-2 hours, embryos were collected and dechorionated in 100% bleach for 1.5 minutes. Embryos were mounted in a cylindrical low-melt agarose gel with fluorescent beads and placed in a custom-built light sheet microscope for imaging. This setup has been previously described in (Streichan, 2018). Using the acquisition software Micro-Manager (https://micro-manager.org/), images were acquired with a frame rate of 1/15 seconds in 8 angles, 45 degrees apart from each other to cover the entire embryo.

##### Quantitative image analysis for confocal microscopy

Confocal images were exported as .tif files from LAS AF Lite software and were imported into MATLAB for image analysis. Different custom scripts were written for different analyses performed. Using ilastik (Version 1.3.3), a machine-learning based segmentation software, probability matrix was generated and exported. Pixel Classification pipeline was performed with these features: Gaussian Smoothing, Laplacian of Gaussian, Gaussian Gradient Magnitude, Difference of Gaussians, Structure Tensor Eigenvalues, Hessian of Gaussian Eigenvalues. For confocal images, Sigma 0.30, 0.70, 1.00, 1.60, 3.50, 5.00, and 10.00 were used for all mentioned features. For light sheet images, Sigma 5.00, 10.00, and 20.00 were used. This probability matrix was imported into MATLAB to generate nuclear masks. 2D convex hull was defined computationally, and the distance from nuclei on the convex hull to the embryo edge was approximated as the distance from cortex.

**Quantitative image analysis for light sheet microscopy**
The acquired images from light sheet microscopy were fused and deconvolved by using the multi-view reconstruction pipeline in ImageJ, which was originally described in [3]. The configurations used for defining dataset are followed, otherwise left as default. Type of Dataset: Image Stacks (LOCI Bioformats), Multiple timepoints: YES, Multiple channels: NO, Multiple illumination directions: NO, Multiple angles: YES, Calibration Type: Same voxel-size for all views, Calibration Definition: User define voxel-size(s). The voxel size is given by the resolution 0.2619 $\mu m$ in x and y (which is unique to our setup), and z depth was 1 $\mu m$. The configurations used for detection of interest points are followed, otherwise left as default. Type of interest point detection: Difference-of-Gaussian, Label interest points: beads, Subpixel localization: 3-dimensional quadratic fit, Interest point specification: Interactive. Parameters were adjusted such that good number of beads were correctly identified while only few to no embryo structure were identified as beads. For registration, Fast 3d geometric hashing algorithm was used. Each time points were registered individually or semi-global registration was used with 5 consecutive time points, depending on the quality of the data. Parameters not mentioned were left as defaults. For deconvolution, 2-5 iterations were used depending on the quality of the image, with automatic PSF extraction from beads with manually defined blocksize 1024x1024x768. The computation was performed on NVIDIA Tesla M10. The fused image stacks were fed to ilastik (Version 1.3.3), which gave a probability map of identical size to the image stack, where each voxel contains a probability number of being the segmentation or not (in this case, either nuclei or spindle, depending on the dataset). The output probability map of ilastik was then imported to our own computational pipeline written with MATLAB. To save time and computational memory resource, we performed Fourier transform on the probability map and convolved it with a Mexican hat (Laplacian of Gaussian) filter, which let us detect and amplify spherical object signals. We then performed inverse Fourier transform and simple thresholding to acquire final segmentation.

**Analysis of migration curves**
Each migration curve in Figure 3D was fitted in MATLAB with a logistic custom equation $x(t) = x_0 + L/(1 + e^{-(t-t_0)/\sigma})$ This implies that the curve's starting value is $x_0$, height of the curve is $L$, and $t_0$ is the time when the value is at the sigmoid midpoint. Then, the time derivative of x(t), v(t), is given as $v(t) = (L/\sigma)/(1 + e^{-(t-t_0)/\sigma})^2 e^{-(|t-t_0|)/\sigma}$ . The maximum velocity occurs at $t = t_0$, which is $v_{max} = L/4\sigma$ (Figure 3G). All fits in this manuscript are performed using the Levenberg-Marquardt shooting-and-matching method.

**Estimation of spindle orientation**
All spindles and the vitelline membrane were segmented using ilastik and custom MATLAB scripts. This gave each spindle's eigenvector, namely spindle$_{eg}$. For each spindle, the corresponding closest point on the vitelline membrane was calculated. Next, a circular patch around the closest point was generated using simple distance threshold of 30 voxels ($\sim 7.86\mu m$). Such a curved patch can be approximated as a plane due to its size. The x, y, z coordinates of all voxels in this circular patch were stored as an N by 3 matrix (xyzn). The center of mass ($\vec{X}_{cm}$) for the plane was approximated as mean of xyzn. Then, we ran singular value decomposition of the vector (xyzn - $\vec{X}_{cm}$) to estimate a unit vector normal to the approximated plane, which was used to calculate the tilt.

**Statistical analysis**
All statistical analyses were performed using JMP Pro. Statistical comparison between two experiments was performed by two-sample t-test. Statistical comparisons between multiple experiments were performed by one-way ANOVA followed by Tukey's test to compare all pairs.

**Analysis of single nuclei RNAseq data** [Albright et al., 2022]

Single nuclei RNAseq data from CITE was analyzed using the available pipeline from the Google Colab notebook in the GitHub repository: https://github.com/aralbright/2021_AAMSME. Specifically, the analysis used the "scVI_clustering.ipynb" notebook with input data "adata_b4batchcorrect.h5ad". The scVI model was retrained during the analysis. Once the initial clustering was obtained, we modified the pipeline to retain the yolk and pole cell nuclei while removing low quality nuclei. Re-clustering was then performed on these filtered nuclei following the pipeline for differential expression analysis.

**Morphometric analysis**

To perform the morphometric analysis we extract a mesh manifold using an in-house open crust algorithm written in Matlab 2021b built upon the openly accessible code by Giaccari Luigi (2009, Matlab file exchange) called Surface Reconstruction from scattered points cloud (open surfaces)(https://www.mathworks.com/matlabcentral/fileexchange/63731-surface-reconstruction-from-scattered-points-cloud-open-surfaces). A kink criteria (>4 curvature over 5 points) is used to threshold for spurious defects and reject vertices. The delaunay mesh is then used to define a dual Voronoi mesh and all geometric quantities such as area, topology, perimeter are calculated in a standard manner from the Voronoi mesh. Calculation of the distance of nuclei from the cortex is based on computing the boundary of the entire nuclear population using the convex hull.

**Simulation of expansion by repulsion**

We initialize the system with 50 particles distributed equidistally on an ellipse of eccentricity $\epsilon$. The average distance is then calculated and saved as $\langle d \rangle$. The initial distance is defined as $d_0$. The force is calculated from the aster-aster repulsion model (presented in supplementary note) with a maximal length $l_m = 1.5\langle d \rangle$. The system is then dynamically relaxed by moving the points according to $\dot{x}_i = \sum_j f_{ij}$. After relaxation spanning timescale $t_{cycle}$, the number of particles is doubled, eccentricity of the ellipse is measured and the entire process is repeated until eccentricity converges. The interaction with the boundary is captured by a steep harmonic potential with a cutoff at $d_0/5$, so the particles only interact with boundary when they are sufficiently close.

**Simulation to estimate torque from neighbours**

We define an n-sided polygon by choosing n-points radially $R(1 + 0.1 * URAND)$ distance away from center and azimuthally $(2\pi/n)(1+0.1*URAND)$ where $URAND$ is an uniform random variable $\in [0, 1]$. Area and perimeter is estimated and saved. We perform an integration of $dl/R(l)$ for each edge segment connecting vertices i and j defined as $\vec{L}_{ij} = \vec{X}_j - \vec{X}_i$. We choose a fraction $p$ of the edge $\vec{L}_{ij}$ by drawing a random variable $p \in [0, 1]$, then we perform a numerical integration on the grid of 1000 points along $\vec{L}_{ij}$ to estimate $\sum_{k=1}^{1000} dl_k/R_k$. The step is then repeated for all edges.

### Mechanics of aster-aster interactions

#### Pushing forces leading to expansion

To capture the mechanics of the aster-aster interactions we consider a simple geometric setup presented in Fig.3A. The aperture of the overlap $\phi_m$ is set by the maximal length of microtubules $l_m$ and the distance $d$ between the aster conics and is given by $\phi_m = \cos^{-1}(d/2l_m) = \cos^{-1}(x)$. The total pushing force experienced by the asters in this 2D analog can be calculated across the whole aperture. If the magnitude of the force exerted along the microtubules is saturated at a value of $f_0$ then the force balance is given by

$$\gamma_t \dot{x} = F_{\text{pushing}} = 2f_0\sqrt{1 - x^2} = -\frac{dW}{dx} \quad , \tag{1}$$

and can be presented as a relaxation phenomena in a potential function $W$ (see Fig.3B). Alternatively one can consider a model where the force generated at each microtubule pair is proportional to the overlap length, hence maximal at the equator and minimal at the periphery. In this case the force balance would be modified to

$$\tau \dot{x} = F_{\text{pushing}} = 2f_0\sqrt{1-x^2} - 2f_0 x \cos^{-1}(x) \quad . \tag{2}$$

#### Alignment torque between neighbouring asters

If the aster overlap is completely symmetric across the equator of the overlap region, the orthogonal component $f_y$ (Fig.3A) completely cancels out due to symmetry. Now if we introduce a small tilt $\psi$ as shown in Fig.4A, the overlap region shrinks by an aperture of $\psi$ below the equator. As a result a clockwise torque is generated and the dynamical equation of tilt $\psi$ is given by

$$\gamma_r \dot{\psi} = -T_0 \int_{-\phi_m+\psi}^{\phi_m} \tan(\phi) d\phi = \tau_0(\psi) \ , \tag{3}$$

where $T_0 = f_0 d/2$ is a maximal torque and $\gamma_r$ is a rotational friction. The fixed points and stability can be found by using a phase-plane analysis as presented in Fig.4B. $\psi$ has a stable fixed point at $\psi^* = 0°$ and two boundary stable points at $\psi^* = \pm 90°$, separated by unstable fixed points.

For an astral domain the total magnitude of the torque depends on the integrated effect around the periphery to account for geometric effects (Fig. 4C). For radial asters, the density of microtubules $\rho$ at distance $R$ from the center $\rho(R) \sim 1/R^2$ [1], while the unit torque $(\vec{R} \times \vec{f})$ generated by each microtubule contact $\sim R$. Hence, along the periphery of the Voronoi domain for a small arc $dl$ (see Fig.4C), we can define the torque element and total torque as

$$d\tau_m \sim R(l) f_0 w \frac{N_c}{4\pi R(l)^2} p(l) dl = f_0 w \frac{N_c}{4\pi} p(l) \frac{dl}{R(l)} \quad \text{and} \quad \tau_m = f_0 w \frac{N_c}{4\pi} \oint p(l) \frac{dl}{R(l)} \quad , \tag{4}$$

where $f_0$ is a typical force exerted at a microtubule contact, $w$ is the width of the nuclear shell and $N_c$ is the number long microtubule emanating from the centrosome. The parameter $p(l)$ is the probability of antiparallel overlap at the location $l$ around the periphery. The geometric scaling reveals that the torque is scale invariant. In other words, if we consider a nuclear shell where the thickness $w$ remains the same but all the Voronoi domains (and the nuclei-to-nuclei distance as a result) are rescaled by a numerical factor $\lambda$, the torque $d\tau_m \sim d(\lambda l)/\lambda R(l) = dl/R(l)$ is independent of the rescaling factor $\lambda$.

The relation with the topology can be seen in few different ways. For example in a regular regular polygon with $n$ sides, each side makes an angle of $2\pi/n$ at the center and hence the number of sides that fall within the fanning angle $\alpha$ of the aster is $= \lfloor \alpha/n\pi \rfloor$. For each interacting side, the corresponding neighbouring aster can point in the correct configuration with probability $p$. Hence the amount of torque $\tau_m$ is proportional to the number of neighbours $n$ and thereby to the topological charge $q$. An alternative way to derive this relationship is to consider the mean-field case $p(l) = p$ and then we simply estimate $\sum H.dl$. For a polygon with side $n$ the total curvature arises from the overall $2\pi$ (also sum of exterior angles) revolution and the contribution of interior angles at each corner. The sum of the interior angles is $= (n-2)\pi$. Hence the total curvature $= n\pi = (q+6)\pi$, where $q$ is the topological charge. For an arbitrarily shaped domain with $n$ corners (rather than straight polygons) we can make use of a well-known identity in integral geometry

$$\oint H(l).dl = 2\pi\chi + \sum_{i=1}^{n} \alpha_i \tag{5}$$

where $H = 1/R$ is the curvature, $\chi$ is the Euler characteristic and $\alpha$ is the exterior angle as indicated in Fig. 4C. Eq.(5) is the two-dimensional version of the Gauss-Bonnet theorem that shows that integral curvature $\oint H(l).dl$ is a topological invariant. Here however no simple constraint of exterior angle is known. Much like foams, we can argue that corner only emerge in astral domains when three such domain meet and must be force-balanced leading to $\sim \pi/3$ exterior angle. As a result, $\oint H(l).dl \simeq 2\pi + q\pi/3$. Irrespective of the exact geometric model used to derive the relationship, we can write the total torque as

$$\tau_{tot}(\psi) = f_0 w \frac{N_c}{4\pi}(q+q_0) \int_{-\phi_m+\psi}^{\phi_m} \tan(\phi) d\phi \tag{6}$$

To estimate the fraction of spindles with $\psi > 45°$ as a function of topological charge $q$, we first seek a steady state solution to the Fokker Planck equation

$$\frac{\partial P(\psi)}{\partial t} = -\frac{1}{t_\eta}\frac{\partial}{\partial \psi}\left[\tau_{tot}(\psi)P(\psi) - \beta_{eff}^{-1}\frac{\partial P}{\partial \psi}\right] \quad , \tag{7}$$

which yields a solution of the form

$$P_{ss}(\psi) = \frac{1}{Z}\exp\left(-\beta_{eff}\int_0^{\psi} \tau_{tot}(\psi^{'})d\psi^{'}\right) \tag{8}$$

where $Z$ is a normalization factor and $t_\eta$ is a relaxation timescale of the torque due to viscous dissipation. The above integral is estimated numerically. The fraction of spindles tilted beyond 45° can be found as

$$F(\psi > 45°|q) = 1 - \int_{0°}^{45°} P_{ss}(\psi) \quad . \tag{9}$$
